# Carboxymethyl Dextran Coated Iron Oxide Magnetic Nanoparticles as Vehicles for CRISPR Ribonucleoproteins

**DOI:** 10.64898/2026.10.28.762406

**Authors:** Jimena Pérez-Maroto, Carmen Escalona-Noguero, Mónica Dhanjani, Nuria Lafuente-Gómez, Milagros Castellanos, Yurena Luengo, Carmen Mayoral-Campos, Gorka Salas, Mirentxu Santos, Marta García, Ángeles Mencía, Rodolfo Murillas, Álvaro Somoza, Begoña Sot

## Abstract

The efficient and safe intracellular delivery of CRISPR ribonucleoproteins (RNPs) remains a major challenge for the development of genome editing strategies. Magnetic iron oxide nanoparticles (MNPs) offer an attractive and biocompatible delivery platform. However, their use for the delivery of functional CRISPR RNPs remains comparatively underexplored. Here, we developed carboxymethyl dextran (CMD)-coated MNPs as non-viral carriers for Cas9 and Cas12a RNPs. Nanocomplexes were formed through electrostatic interactions, without the need for chemical coupling reagents or additional surface functionalization. Both RNPs efficiently associated with MNPs, and the resulting nanocomplexes exhibited stable physicochemical properties and retained their cargo in serum-supplemented culture medium for up to 24 hours. Cellular uptake was confirmed by microscopy, while biocompatibility assays revealed no overt cytotoxicity. Functional genome editing was demonstrated at reporter and endogenous gene loci in HEK-293T cells, as well as in primary human dermal fibroblasts. Finally, we explored the therapeutic potential of this platform for recessive dystrophic epidermolysis bullosa (RDEB) by targeting COL7A1 exon 80, with the aim of removing the prevalent c.6527insC mutation while preserving the reading frame. Cas9 RNP nanocomplexes enabled exon 80 deletion. These findings establish CMD-coated MNPs as a versatile platform for the delivery of Cas9 and Cas12a RNPs and support further investigation of magnetic nanoparticle-mediated genome editing for inherited skin disorders.

## 1. Introduction

Clustered Regularly Interspaced Short Palindromic Repeats (CRISPR)–associated systems have revolutionized genome editing by enabling precise, efficient, and programmable modification of genetic sequences. Originally derived from prokaryotic adaptive immune mechanisms, CRISPR-Cas systems rely on RNA-guided nucleases capable of inducing site-specific DNA double-strand breaks, which are subsequently repaired by endogenous cellular pathways such as non-homologous end joining (NHEJ) or homology-directed repair (HDR), enabling gene disruption or precise sequence correction ^1^. Recent years have witnessed major achievements in the clinical translation of CRISPR-based technologies, including the approval of ex vivo gene-edited therapies and the advancement of multiple in vivo clinical trials targeting genetic diseases. Despite these advances, significant challenges remain. One of the most critical limitations is the efficient and safe delivery of CRISPR systems, particularly in vivo. Achieving targeted delivery to specific tissues, ensuring intracellular uptake, and minimizing off-target effects and immune responses continue to be major bottlenecks in the clinical application of genome editing technologies^2^ ^3^.

CRISPR components can be delivered in different formats, including plasmid DNA, messenger RNA (mRNA), or ribonucleoprotein (RNP) complexes, consisting of preassembled Cas protein and guide RNA. Among these, RNP delivery offers several advantages. RNPs enable rapid editing upon delivery and exhibit transient activity, thereby reducing the risk of off-target effects, prolonged nuclease exposure, and genotoxicity. In contrast, plasmid-based delivery may lead to sustained expression, increasing the likelihood of undesired edits and insertional mutagenesis. Compared to mRNA-based approaches, RNP delivery eliminates the need for translation, allowing faster onset of action and potentially lower immunogenicity.

The need for improved delivery systems has driven growing interest in nonviral vectors, particularly nanoparticles, as alternatives to viral platforms. While viral vectors such as adeno-associated viruses (AAVs) provide high transduction efficiency, they are associated with limitations including immunogenicity, limited cargo capacity, high production costs, and risks of genomic integration.^4,5^ In contrast, nanoparticles offer tunable physicochemical properties, improved safety profiles, reduced immunogenicity, and the possibility of repeated administration. Nanoparticles have demonstrated significant progress as delivery platforms for nucleic acids and gene editing tools. Various nanomaterials, including lipid nanoparticles, polymeric nanoparticles, gold nanoparticles, and inorganic nanocarriers, have shown the ability to protect cargo from degradation, facilitate cellular uptake, and enable targeted delivery to specific tissues. These systems have been successfully applied in preclinical and clinical settings, particularly for liver-targeted therapies, highlighting their versatility and translational potential.^2,5^ However, most strategies focus on DNA or mRNA delivery rather than RNPs, and the efficient delivery of RNPs remains an important area for further improvement.

Among inorganic nanomaterials, magnetic iron oxide nanoparticles (MNPs), including Fe₂O₃-based magnetic nanoparticles, have attracted considerable attention due to their unique magnetic properties, biocompatibility, and ease of surface functionalization^6^. These nanoparticles can exhibit superparamagnetic behavior, enabling external magnetic field-guided targeting and enhanced accumulation at specific sites^7^. Furthermore, their high surface area allows efficient loading of therapeutic agents, including drugs and nucleic acids, making them promising candidates for gene delivery applications. MNPs have been widely explored in drug delivery, cancer therapy, and imaging, as well as in nucleic acid delivery systems such as plasmid DNA, siRNA, and miRNA^8^. Building on this broad potential, magnetic nanoparticles are now being investigated for CRISPR delivery. To date, most studies involving MNPs have focused on the delivery of plasmid-based CRISPR systems, leveraging electrostatic interactions and magnetic targeting to enhance transfection efficiency^7,9,10^. However, the delivery of CRISPR RNPs using magnetic nanoparticles remains comparatively underexplored, despite its potential advantages in terms of safety and editing precision. Surface modification plays a crucial role in optimizing nanoparticle performance. In this context, coatings such as carboxymethyldextran (CMD) have been extensively used to improve colloidal stability, biocompatibility, and circulation time of magnetic nanoparticles^11,12^. CMD-coated iron oxide nanoparticles have been employed in various biomedical applications, including drug delivery^13^ and imaging^14^, and some formulations have received regulatory approval from the U.S. Food and Drug Administration (FDA)^15^, underscoring their clinical relevance.

In this work, we investigated CMD-coated iron magnetic nanoparticles as carriers for CRISPR ribonucleoprotein (RNP) complexes. Cas9- and Cas12a-based nanocomplexes were characterized in terms of their physicochemical properties, colloidal stability, and biocompatibility in vitro. Their genome-editing activity was evaluated in mammalian cells by targeting the endogenous B2M gene, and the feasibility of applying this approach to a disease-relevant target was further explored by inducing the deletion of exon 80 in COL7A1 in human dermal fibroblasts. Together, these experiments provide an initial assessment of CMD-coated magnetic nanoparticles for RNP delivery and establish a basis for further studies aimed at improving nonviral CRISPR delivery.

## 2. Results

### 2.1. Assembly, physicochemical characterization, and stability of MNP–RNP nanocomplexes

Electrostatic interaction between charged MNPs and cargo has been succesfully used for non viral delivery in vitro and in vivo^7,9,10^. The simplicity of attachment, avoiding chemical corsslinking, favours the scalability of the nanocomplex production. Fe_2_O_3_- based magnetic nanoparticles were prepared by coprecipitation, leading to an almost spherical morphology core with a mean size of 14 nm (Figure S1) that was successively coated with CMD. Thus, MNP-RNP nanocomplexes were produced by electrostatic interaction incubating CMD-coated MNPs, negatively charged, with Cas9 or Cas12 RNPs in low salt buffer (binding buffer). Although Cas proteins have a negative surface in complex with a guide RNA, more pronounced in the case of Cas9 due to the longer length of its gRNA, they also present positive surfaces able to interact with the negative charges of the CMD-coated MNPs, including NLS sequences added for nuclear internalization (Figure S2). MNPs and RNPs were mixed in binding buffer and incubated overnight at 4°C. The binding efficiency was analyzed by co-precipitation and SDS-PAGE of supernatant and resuspended pellet samples. The percentage of MNP-bound protein was determined by gel densitometry (Figure S3). This analysis confirmed that both RNPs efficiently bound to the MNP-CMD surface, reaching binding capacities of 500-600 pmol of RNP per milligram of iron, corresponding to a binding efficiency of 60-70% (Figure 1A). The binding was slightly higher for Cas12a RNPs, probably due to the above-mentioned differences in the surface charge of Cas9 and Cas12a RNPs.

**Figure 1.**
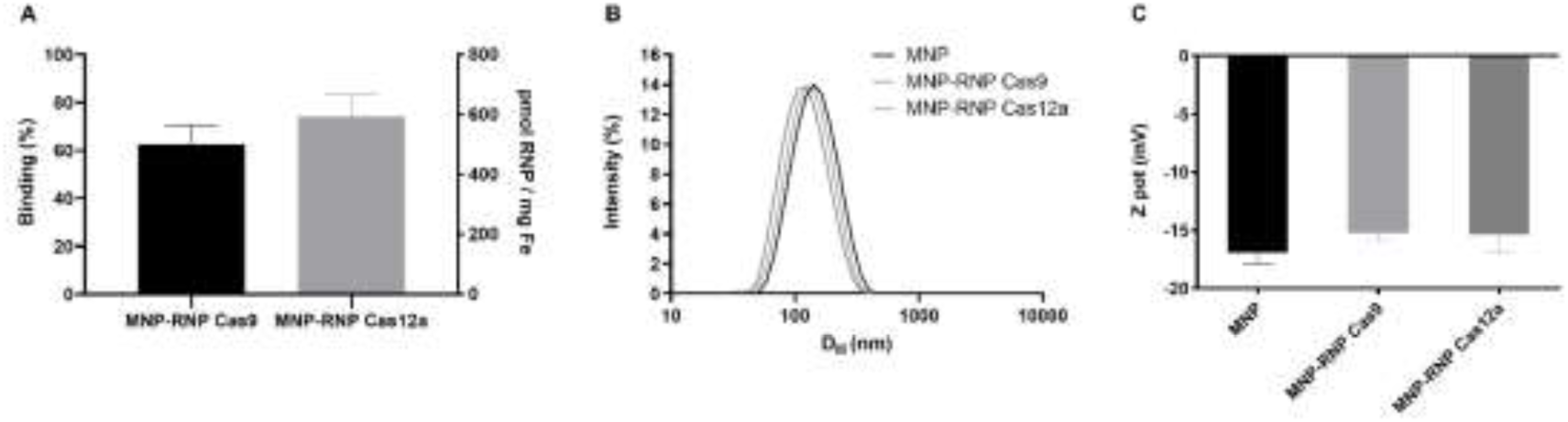
Characterization of the nanocomplexes. A) Binding of Cas9 and Cas12 RNPs to MNPs measured by co-precipitation followed by sds-page and densitometry analysis. The binding is expressed as % of RNP bound to the MNP and pmol RNP per mg of Fe. B) MNP and MNP-Cas nanocomplex size (hydrodynamic diameter) determined by DLS. C) Z-potential measurements of MNPs and MNP-Cas nanocomplexes.

MNP-RNP nanocomplexes were further characterized using DLS to determine their size (hydrodynamic radius) and surface charge (zeta potential). The size of the nanocomplexes was similar, and even slightly smaller, to that of the control without the protein, meaning that the binding with RNPs did not destabilize the MNPs (Figure 1B) (Table 1). While CMD-coated MNPs are highly negative, zeta potential became less negative for MNP-RNP nanoconjugates, further confirming that Cas RNPs are interacting with the NP surface (Figure 1C). Altogether, these observations confirmed the formation of stable MNP-RNP nanoconjugates. Both Cas9 and Cas12 can form nanocomplexes with MNPs, showing the versatility of our strategy.

**Table 1.** Nanocomplexes colloidal characterization. Hydrodynamic diameter and polydispersity index (PDI) of MNP and MNP-Cas RNP nanocomplexes. (N=3)

| Sample | Size (nm) ± SD | PDI ± SD |
| --- | --- | --- |
| MNP | 130.7 ± 1.42 | 0.15 ± 0.02 |
| MNP-RNP Cas9 | 120.3 ± 1.4 | 0.14 ± 0.01 |
| MNP-RNP Cas12a | 111 ± 1.7 | 0.13 ± 0.01 |

The development of nanosystems for biomedical applications requires assessing their behavior in biological media. Upon contact with biological fluids, proteins and other biomolecules rapidly adsorb onto the MNPs’ surface, forming the protein corona. This coating can affect their physicochemical properties -such as size and superficial charge-affecting colloidal stability and subsequent cellular interactions^16,17,18^. The impact of protein corona formation on MNP-RNPs’ colloidal stability was evaluated by DLS after 0, 4 and 24 hours of incubation in DMEM supplemented with 10% FBS, the most common condition used in vitro. Incubation of uncomplexed CMD-MNPs with the protein rich medium promoted their aggregation, potentially due to protein adsorption onto their negatively charged surface. Their hydrodynamic diameter reached their maximum after 4h of incubation, followed by stabilization at 24h (Figure 2A; Table S1). The DLS profiles and the PDI measurements indicated the presence of two subpopulations with different degrees of aggregation. In contrast, both Cas9 and Cas12a MNP-RNPs exhibited high colloidal stability, with only minimal increases in hydrodynamic diameter and PDI over time (Figure 2A; Table S1). These findings suggest that RNP complexation modifies the interactions between the MNP surface and serum proteins, potentially influencing protein corona formation and contributing to the enhanced colloidal stability of the nanocomplexes (Figure 2A; Table S1).

**Figure 2.**
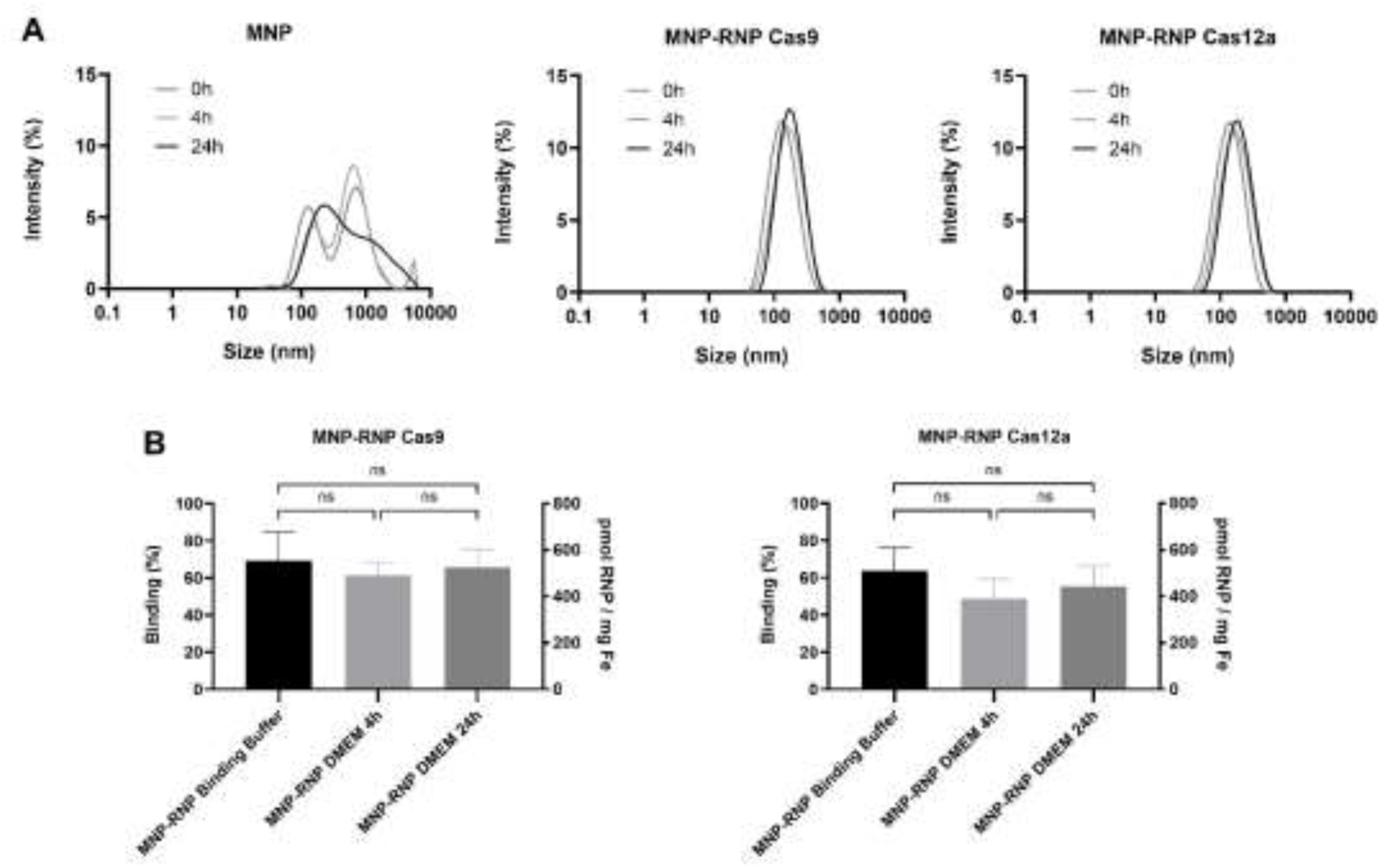
Nanocomplexes characterization in biological media. A) Hydrodynamic diameter of MNP and MNP-Cas RNP nanocomplexes (N=3) in 10% FBS at 4h and 24h. B) RNP bound to MNPs after incubation with 10%FBS.

A critical requirement for MNP-based delivery systems is the prevention of premature cargo release upon exposure to biological fluids. To evaluate whether protein corona formation compromises nanocomplex integrity, we assesed RNP retention on the MNP surface following incubation in DMEM supplemented with 10% FBS. No significant displacement of either Cas9 or Cas12a RNPs was observed (Figure S4), and the nanocomplexes maintained a binding efficiency of approximately 60% for up to 24 hours (Figure 2B). These findings indicate that the interaction between the MNPs and RNPs is sufficiently strong to support their use as effective gene editing delivery carriers.

### 2.2. Biocompatibility of MNP–RNP nanocomplexes

Assessing the cytotoxicity of MNP-RNP nanocomplexes is essential for evaluating their translational potential. Accordingly, we determined cell death in HEK-293T cells after 24 hours of incubation with MNP–RNPs using Annexin V and Propidium Iodide staining followed by flow cytometry. To assess the toxicity of the biomaterial independently of protein activity, RNPs were formed using guide RNAs (AAVS1, Table S2) with limited activity (Figure S5). Apoptosis and necrosis were monitored at 24, 48, and 72 hours to characterize the long-term cellular response. MNPs alone and MNP-RNP Cas12a did not induce cytotoxicity, showing a high degree of biocompatibility (Figure 3A, Figure S6). MNP-RNP Cas9 treatment produced a slight increase in the proportion of early apoptotic cells during the first 24 hours of treatment (Figure 3A, Figure S6). This proportion remained stable over time and decreased by 72 hours post-treatment (Figure 3A).

**Figure 3.**
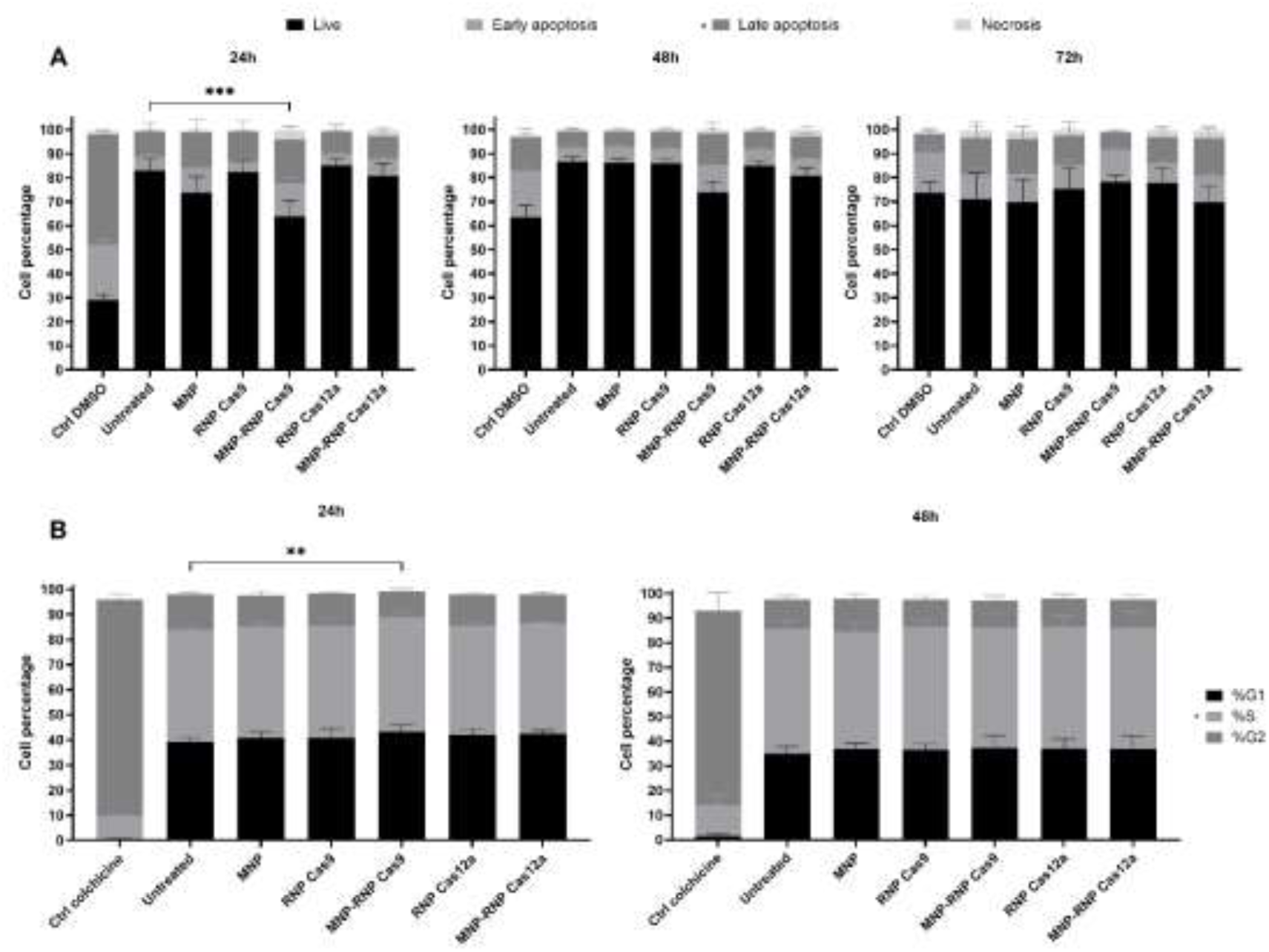
Nanocomplexes biocompatibility. A) Cell death in HEK-293T cells after 24 hours of incubation with MNP–RNPs using Annexin V and Propidium Iodide staining followed by flow cytometry, at 24, 48 and 72h. B) Cell-cycle determination after treatment with MNP-RNP complexes. Values represents the mean ±SD of three replicates at 24h and 48h. Statistical analysis was performed using one-way ANOVA test (each group vs. untreated) ** P<0.01, ***P<0.001.

Although the increased proportion of apoptotic cells suggested that the treatment may induce a measurable cellular stress response under the experimental conditions tested, the proportion of live cells decreased from 85% in untreated controls to 65% in treated cells, a relative reduction of 23.5%. This value is below the 30% reduction threshold described in ISO 10993-5:2009 as indicative of cytotoxicity.

Cell cycle progression was also assessed to determine whether nanocomplex internalization affected cellular homeostasis. Following treatment, HEK-293T cells were stained with propidium iodide and analyzed by flow cytometry at 24- and 48-hours post-treatment, to monitor the kinetics of the cellular response. MNP–RNP Cas9 treatment induced a small but statistically significant increase in the proportion of cells in S phase at 24h, whereas the slight changes observed in the G1 and G2 phases were not statistically significant (Figure S7). This effect was no longer observed at 48 hours (Figure 3B). This result was consistent with the cell-death results (Figure 3A), indicating a limited effect on cell-cycle distribution, although the underlying mechanism remains to be elucidated.

### 2.3. Cellular uptake and intracellular delivery of MNP–RNP nanocomplexes

The overall therapeutic effectiveness of MNP-RNPs relies on their ability to interact with the cell surface and trigger cellular uptake. To evaluate these particle-cell interactions, HeLa cells were incubated with MNP-RNP nanocomplexes and stained with Prussian blue, a reagent commonly used to visualize iron. MNP-RNPs remained associated with the cells even after several 1X PBS washing steps, confirming successful surface binding and initial internalization (Figure 4A). To further characterize cellular uptake, nanocomplexes were assembled using fluorescently labeled RNPs (ATTO-655). Following a 3-hour incubation, live-cell fluorescence imaging confirmed successful uptake for both Cas9 and Cas12a RNPs. The ATTO signal showed a punctate pattern predominantly located in the cytosol, with additional signal detected in the nucleus, where gene editing occurs (Figure 4B). Altogether, these findings confirm that MNPs are suitable non-viral vectors for the cellular uptake and intracellular delivery of CRISPR RNPs.

**Figure 4.**
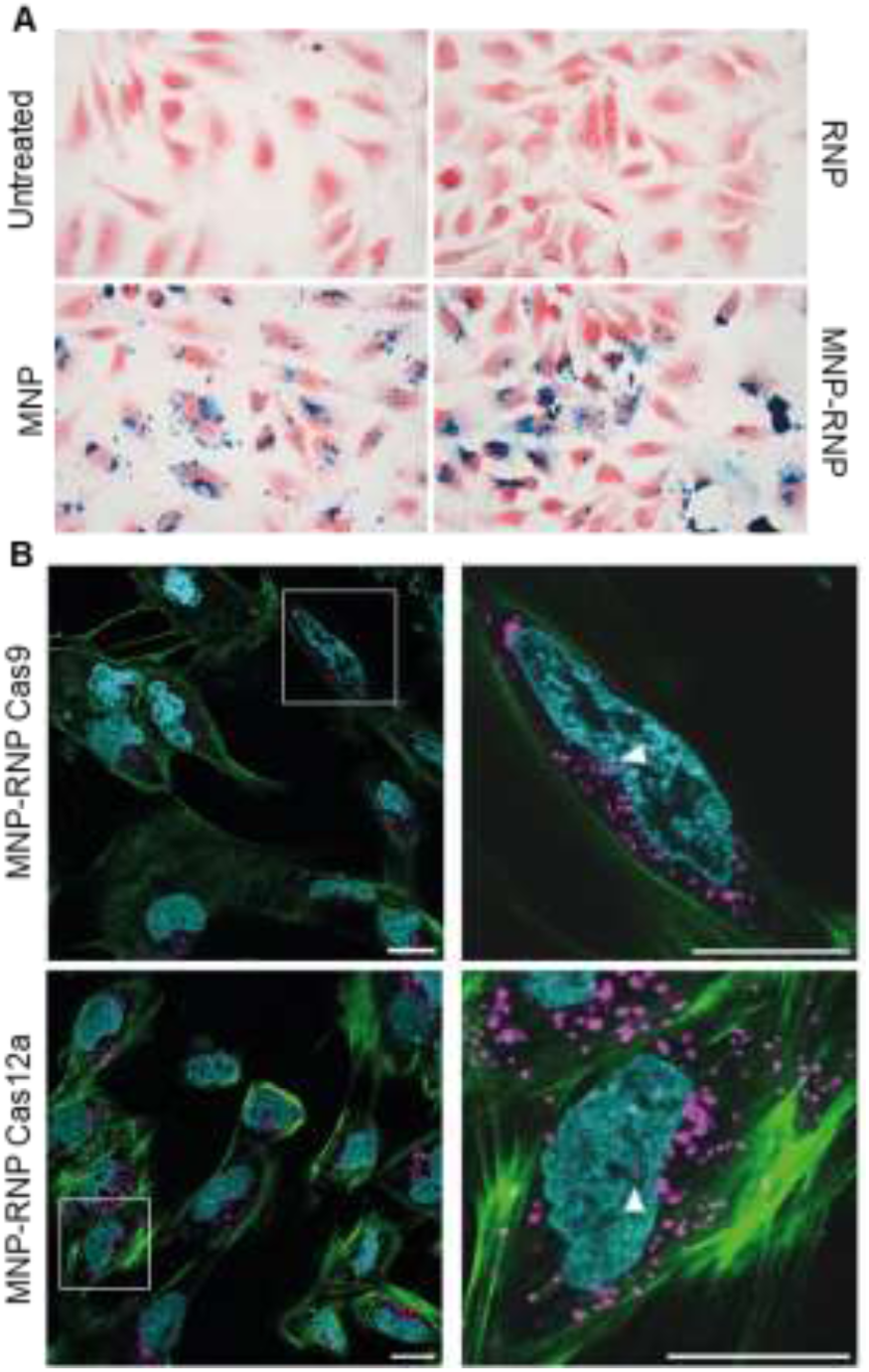
Nanocomplexes uptake by HeLa cells. A) wide-field microscopy images of HeLa cells treated with MNP-RNP complexes and stained with prusian blue for Fe visualization. B) Live cell confocal images of HeLa cells treated with MNP-RNP nanocomplexes. RNPs labelled with ATTO (magenta), nuclei by Hoeschst (cyan) and actin by SPY555 (green). Scale bar: 20µm.

### 2.4. Optimization of MNP–RNP-mediated gene editing in reporter cell lines

Altogether, these findings indicate that MNPs are suitable vehicles for the intracellular delivery of CRISPR RNPs. We next evaluated gene editing activity of the MNP-RNP nanocomplexes using fluorescence-based reporter cell lines. First, MEF-tdTomato cells were used to optimize the working concentration. In this cell model, tdTomato gene expression is repressed by an upstream frt-flanked STOP signal. Designing a Cas9 guide RNA that targets both frt sites enables excision of the STOP signal, thereby restoring the expression of the fluorescent tdTomato protein^19^ (Figure 5A). Cells were treated for 4 h with Cas9 RNP concentrations of 200 or 400 nM, corresponding to the amount of RNP bound to MNP–RNP nanocomplexes, assuming a binding efficiency of 50%. Therefore, the total RNP concentration added to the cells was 400 or 800 nM, respectively. RNP was delivered using MNP–RNP nanocomplexes or Lipofectamine 2000. Gene editing efficiency was markedly higher at 400 nM than at 200 nM. Furthermore, at 400 nM the efficiency was comparable to that achieved by lipofection (Figure 5A). Hence, an RNP concentration of 400 nM of RNP was used in subsequent experiments.

**Figure 5.**
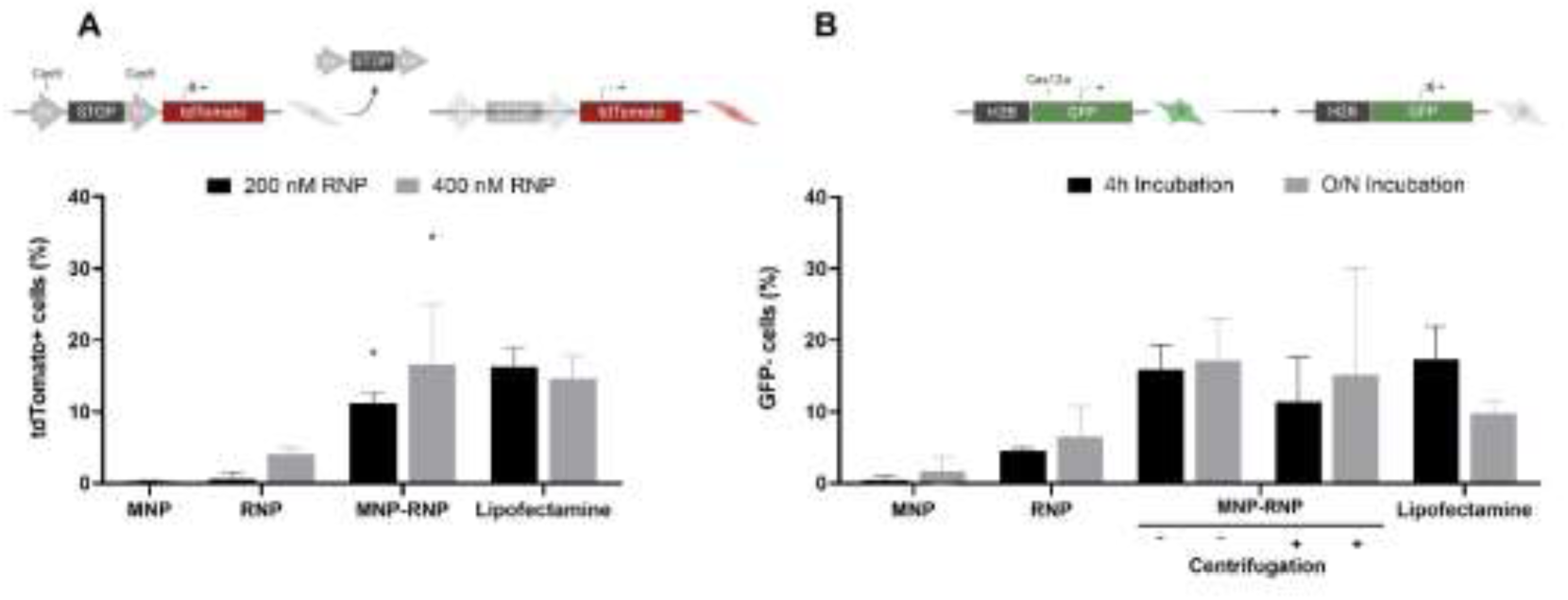
Nanocomplexes gene editing optimization. A) Cas9 gene editing efficiency measured by percentage of tomato positive cells by flow cytometry. B) Cas12a gene editing efficiency measured by percentage of GFP positive cells by flow cytometry. Statistical analysis was performed using two-way ANOVA test. * P<0.05

The HEK-293-GFP cell line, which constitutively express a histone H2B-GFP fusion protein, was selected to assess the activity of the MNP-RNP Cas12a nanocomplexes. To evaluate CRISPR-mediated knockout efficiency, cells were treated with the nanocomplexes containing Cas12 RNPs with a crRNA targeting the GFP gene. Flow cytometry was used to quantify the reduction in GFP expression as a functional readout of gene-editing activity. Our previous results (Figure 1A) showed that approximately 60– 70% of the protein added during nanocomplex formation binds to the MNPs, indicating that the resulting preparations contain a fraction of unbound RNPs. To determine whether this free RNP fraction affected the gene-editing activity of the nanocomplexes and whether its removal was necessary, nanocomplexes were either used directly or subjected to centrifugation to remove unbound protein before cell treatment. The percentage of gene knock out achieved with centrifuged MNP-RNPs was comparable to that obtained with non-centrifuged complexes (Figure 5B), suggesting that the unbound RNP fraction did not significantly contribute to gene editing and supporting the ability of MNPs to act effective carriers for Cas RNPs. Furthermore, the centrifugation step increased the variability of the results, and therefore this step was omitted from the final protocol. We also investigated whether the incubation time used for nanocomplexes formation affected transfection efficiency. Although lipofection efficiency decreased when RNPs were incubated overnight, no reduction was observed for the MNP-RNPs (Figure 5B). These findings suggested that MNPs may help preserve RNP activity during nanocomplex formation. No differences in RNP binding (Figure S8) or gene editing efficiency (Figure 5B) were observed between the two incubation times.

### 2.5. Functional genome editing in cell lines and primary human dermal fibroblasts

To evaluate the efficacy of MNP-RNP nanocomplexes for editing endogenous genes, we first targeted the endogenous beta-2-microglobulin (B2M) gene. B2M encodes an essential light chain of the Major Histocompatibility Complex class I (MHC-I), which is required for its cell-surface expression. Thus, successful B2M knockout directly leads to the loss of cell-surface MHC-I, enabling straightforward quantification by flow cytometry. Following overnight incubation of HEK-293T cells with MNP-RNP Cas9 or MNP-RNP Cas12a, genome editing was assessed 5 days post-transfection using ddPCR to quantify gene-editing and flow cytometry to measure the reduction in B2M expression. Both formulations produced non-homologous end joining (NHEJ) measured by ddPCR, with indel frequencies of ∼14% for both Cas9 and Cas12a MNP-RNP complexes (Figure 6A,B). This gene editing efficacy was associated with a significant increase in the proportion of B2M-negative cells, as determined by flow cytometry (Figure 6C, D).

**Figure 6.**
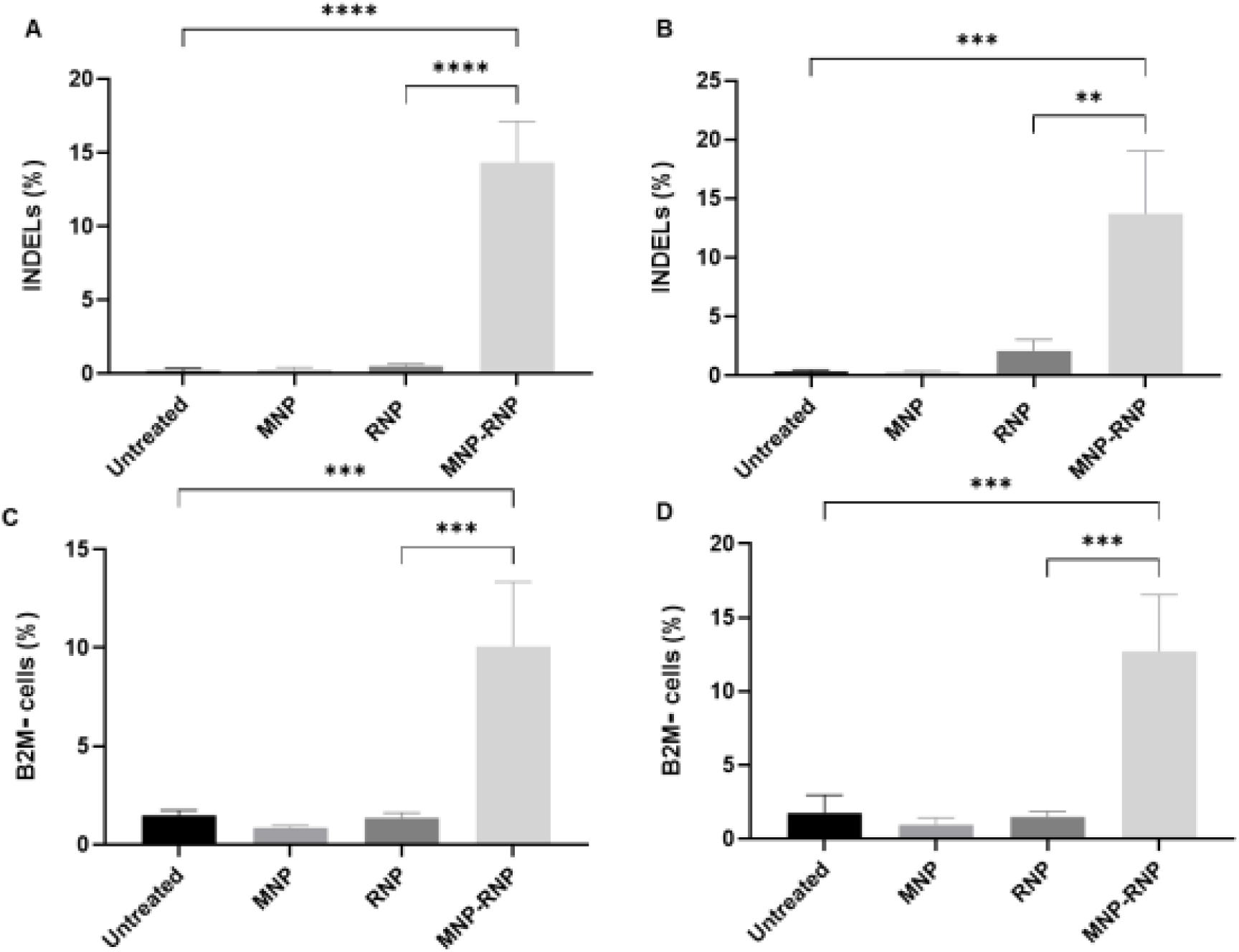
Nanocomplexes gene editing of B2M gene in HEK-293T cells. A) Cas9 gene editing efficiency measured by ddPCR. B) Cas12a gene editing efficiency measured by ddPCR. C) Cas9 gene editing efficiency measured by flow cytometry. B) Cas12a gene editing efficiency measured by flow cytometry. Statistical analysis was performed using one-way ANOVA test. ***P<0.001, **** P<0.0001

Our previous results showed that the RNPs remained bound to the MNPs in biological media. In line with that, we also confirmed that incubation in biological media and the formation of a protein corona did not affect the the nuclease activity of the RNPs inside cells for the knockout of the B2M gene (Figure S9).

The ability of MNP-RNP nanocomplexes to edit the B2M gene in primary cells, was also investigated. Primary human dermal fibroblast from neonatal foreskin (HDFn cells) were treated with Cas9- or Cas12a-based MNP-RNPs containing a guide RNA targeting B2M, and cell-surface B2M expression was assessed by flow cytometry (Figure 7A). Although the results showed greater variability than those obtained in established cell lines, treatment with both nanocomplex formulations resulted in a clear reduction in B2M expression, consistent with successful gene editing.

**Figure 7.**
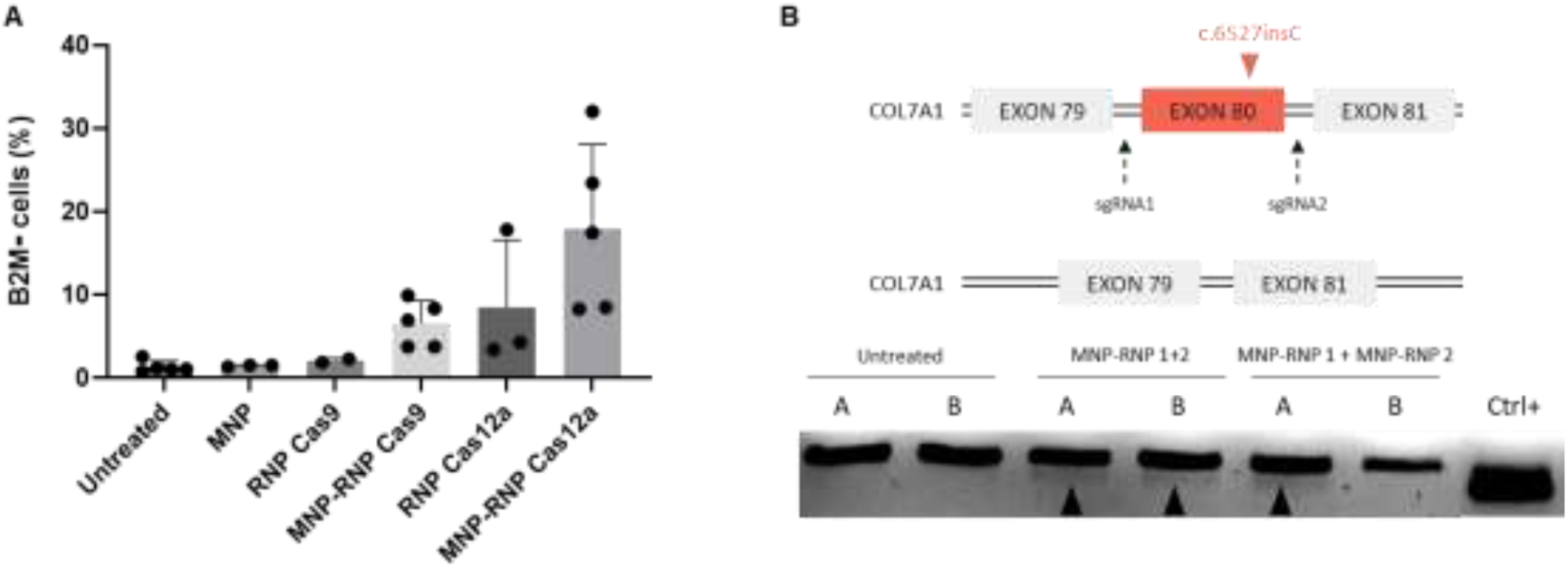
Nanocomplexes gene editing of target genes in HDFn cells. A) Cas9 and Cas12a gene editing efficiency of B2M gene measured by flow cytometry. B) Scheme of gene editing strategy (upper panel) and gene editing of col80 visualized by the appearance of a shorter amplicon of COL7A1.

These promising results prompted us to investigate the potential of our nanocomplexes for gene editing of a therapeutic target in recessive dystrophic epidermolysis bullosa (RDEB). RDEB is a severe inherited skin disorder caused by mutations in COL7A1, the gene encoding type VII collagen, a key structural component of the anchoring fibrils that attach the epidermis to the dermis. Loss or dysfunction of type VII collagen results in extreme skin fragility, chronic wounds, scarring, and a high risk of aggressive squamous cell carcinoma, profoundly affecting quality of life and life expectancy^20^. We had already developed a gene-editing strategy for RDEB based on CRISPR/Cas9-mediated deletion of COL7A1 exon 80, targeting the prevalent c.6527insC mutation in the Spanish RDEB population. This approach uses two distinct CRISPR/Cas9 ribonucleoprotein (RNP) complexes targeting intronic regions flanking exon 80, resulting in its excision (Figure 7B). The resulting in-frame transcript encodes a shorter but functional version of type VII collagen^21,22^. HDFn cells were treated with Cas9-based MNP-RNP nanocomplexes targeting the two intronic regions flanking COL7A1 exon 80. Two formulation strategies were evaluated. In the first, each RNP was independently complexed with MNPs, and the resulting nanocomplexes were mixed at a 1:1 RNP molar ratio. In the second, the two RNPs were mixed at a 1:1 molar ratio before the addition of MNPs to form the nanocomplexes. Genomic DNA was isolated from treated cells, and exon 80 deletion was assessed by PCR followed by agarose gel electrophoresis (Figure 7B). Both formulation strategies resulted in 8-10% exon 80 deletion, measured by gel densitometry. Mixing RNP1 and RNP2 before nanocomplex formation appeared to yield more reproducible results.

## 3. Discussion

Iron oxide magnetic nanoparticles have previously been explored as delivery vehicles for CRISPR components, although most studies primarily focused on plasmid-based systems rather than preassembled RNPs^7,10^. The use of CRISPR RNP complexes instead of plasmid DNA offers important safety advantages, as it eliminates the risk of genomic integration of the plasmid DNA and may reduce the likelihood of off-target editing due to the transient activity of the nuclease. However, the use of MNPs for intracellular protein delivery remains relatively underexplored. To date, the functionalization of MNPs with proteins has primarily been investigated to improve nanoparticle stability or to immobilize targeting ligands, particularly antibodies, rather than to deliver functional protein cargo into cells. More recently, magnetic nanoparticle-mediated delivery of Cas9 RNPs has also been reported^23,24^. In this context, our study extends the application of magnetic iron oxide nanoparticles by demonstrating that CMD-coated MNPs can associate with preassembled Cas9 and Cas12a RNPs through a straightforward formulation procedure and mediate functional genome editing in mammalian cells. Unlike approaches requiring more elaborate carrier architectures^24^, this strategy relies on conventional iron oxide nanoparticles and a simple RNP-loading procedure. Furthermore, unlike the approach reported by Go et al.^23^, in which the formation of MNP–RNP nanocomplexes was not experimentally demonstrated, we directly characterized the association between MNPs and RNPs. By demonstrating the formation of MNP–RNP nanocomplexes, our study provides a stronger basis for further investigating this platform as a potential in vivo delivery vehicle. It is also worth highlighting the simplicity and versatility of nanocomplex formation. This approach is compatible with both of the most widely used CRISPR systems, Cas9 and Cas12a, and relies on electrostatic interactions, allowing MNP–RNP nanocomplexes to form simply by incubation, without the need for additional coupling reagents or complex functionalization procedures. Notably, the strength of these electrostatic interactions appears sufficient to maintain cargo association and intracellular activity under biological conditions, supporting the potential of this platform for in vivo applications. Interestingly, the presence of RNPs on the MNP surface also appears to prevent aggregation in biological media. Although further experiments are required to elucidate the underlying mechanism, these findings suggest that the adsorbed RNPs may act as a relatively stable protein layer, potentially limiting the adsorption and dynamic exchange of serum proteins at the MNP surface and thereby reducing aggregation. This hypothesis is consistent with the concept of a hard protein corona, which describes a layer of proteins that remains tightly associated with nanoparticle surfaces^17^.

A prerequisite for the use of nanocomplexes as intracellular delivery vehicles is their biocompatibility. Our cell death and cell-cycle analyses indicate that neither nanocomplex formulation is overtly cytotoxic in the cell lines tested. Interestingly, MNP– RNP complexes assembled with Cas9 induced a small but statistically significant increase in the proportion of cells in S phase, together with a slight increase in early apoptosis, whereas the corresponding Cas12-based complexes did not produce detectable changes in cell-cycle distribution or apoptosis. Since both formulations contained the same magnetic nanoparticles and exhibited comparable colloidal stability, as assessed by dynamic light scattering (DLS), these differences are unlikely to be explained solely by nanoparticle composition or gross differences in colloidal stability. Furthermore, both systems achieved similar gene-editing efficiencies, suggesting that the two RNP formulations were similarly uptaken by cells. The distinct cellular responses may instead be related, at least in part, to differences in the physicochemical properties of the assembled complexes. Cas9- and Cas12-containing complexes showed different zeta potential. Nevertheless, it reflects the overall electrokinetic properties of the particles and does not resolve local surface charge distributions. Differences in protein surface charge patterns, protein–membrane interactions or intracellular responses to the respective RNP components could therefore contribute to the observed effects. Further studies, including quantitative assessment of cellular uptake and intracellular trafficking, would be required to establish the mechanisms underlying the differential cellular responses to MNP–Cas9 and MNP–Cas12 complexes.

Gene editing was successfully achieved using both Cas9- and Cas12-based nanocomplexes, enabling the modification of both reporter and endogenous genes. Importantly, editing activity was demonstrated not only in established cell lines but also in primary fibroblasts, highlighting the versatility of these nanocomplexes across different cell types and their potential for applications requiring gene editing in clinically relevant primary cells. In this regard, the obtention of gene editing of COL7A1is a promising result for the use of these nanocomplexes in RDEB. Although the extent of obtained gene editing is limited, preclinical studies have shown that partial restoration of type VII collagen (COL7) can improve dermal–epidermal adhesion and skin mechanical integrity in RDEB models. However, the minimum COL7 level required for functional rescue remains uncertain and may depend on the model and duration of follow-up^25,26^.

Finally, the successful delivery of CRISPR/Cas9 and Cas12 RNPs using magnetic nanoparticles to primary fibroblasts opens promising opportunities for their use in in vivo therapeutic gene editing in genodermatoses such as RDEB. Nanoparticle-based delivery systems are attracting increasing interest for the treatment of genodermatoses, including epidermolysis bullosa (EB), as they offer the potential for non-viral delivery of therapeutic nucleic acids and gene-editing tools to affected skin tissues^27–29^. In this context, magnetofection represents a promising approach in which magnetic nanoparticles carrying therapeutic cargo are guided towards a target site by applying an external magnetic field, potentially enhancing local delivery and cellular uptake^30^. This strategy may be particularly advantageous for genodermatoses, in which lesions are often accessible and spatially localized, allowing magnetic fields to be applied directly to selected skin areas. The combination of magnetic nanoparticles with Cas9- or Cas12-based RNPs could therefore provide a versatile platform for localized in vivo gene editing, potentially reducing systemic exposure while enabling repeated treatment of affected regions. Although further studies are required to establish delivery efficiency, tissue specificity, safety and therapeutic efficacy in vivo, this approach offers an attractive avenue for developing non-viral gene-editing therapies for EB and other inherited skin disorders.

## 4. Materials and Methods

### 4.1. AsCas12a and SpCas9 expression and purification

AsCas12a was expressed and purified according to the method described by Moreno-Mateos et al.^31^, with minor modifications. The AsCas12a-2NLS bacterial expression plasmid was introduced into competent *E. coli* BL21 cells. The AsCpf1-2NLS plasmid was kindly provided by Jennifer Doudna (Addgene plasmid #102565). A starter culture was prepared by growing the transformed cells overnight at 37 °C in LB medium supplemented with ampicillin. Subsequently, 75 mL of the starter culture were used to inoculate 3 L of LB-ampicillin medium, which was incubated at 37 °C and 180 rpm until the OD600 reached 0.3–0.4. The incubation temperature was then reduced to 18 °C, and protein expression was induced by adding IPTG to a final concentration of 0.5 mM when the OD600 reached 0.5–0.6. Following overnight incubation, the cells were harvested by centrifugation, and the resulting pellet was resuspended in lysis buffer consisting of 50 mM Tris-Cl pH 7.5, 500 mM NaCl, 10% glycerol, 5 mM β-mercaptoethanol, 0.5 mM PMSF, 0.2 mg/mL lysozyme (Alfa Aesar), DNase (Panreac), and EDTA-free protease inhibitors (Pierce, Thermo Fisher Scientific). The cell suspension was incubated for 1 h at 4 °C and subsequently subjected to sonication to achieve complete cell lysis (3 cycles of 10 s ON and 10 s OFF, at 50% amplitude; Sonic Dismembrator Model 120, Fisherbrand). The resulting lysate was clarified by centrifugation at 10,000 × g for 1 h at 4 °C. Soluble AsCas12a was purified by immobilized metal affinity chromatography using a 5 mL HisTrap HP column (Cytiva). After sample loading, the column was washed with at least 5 column volumes (CVs) of buffer A, containing 25 mM Tris pH 7.5, 0.6 M NaCl, 5 mM MgCl2, 10% glycerol, 10 mM imidazole, and 5 mM β-mercaptoethanol. Protein elution was performed using a combination of an isocratic step with 10% buffer B (50 mM sodium phosphate pH 7.4, 500 mM NaCl, 500 mM imidazole, and 10% glycerol) and a gradient ranging from 10% to 100% buffer B. The protein content of selected chromatography fractions was assessed by SDS-PAGE using 10% acrylamide gels. Fractions containing AsCas12a were pooled and concentrated by centrifugation using a 100 kDa Amicon filter (Merck Millipore), according to the manufacturer’s instructions. The 6His-MBP tag was subsequently removed by cleavage with TEV protease, using 1 mg of protease per 50 mg of total protein. The sample was then dialyzed overnight at 4 °C against dialysis buffer containing 25 mM Tris-Cl pH 7.0, 600 mM NaCl, 10% glycerol, and 1 mM DTT. Following dialysis, the protein solution was centrifuged at 16,000 × g for 15 min at 4 °C to remove potential aggregates. The supernatant was then loaded onto a Superdex 200 size-exclusion chromatography column (GE Healthcare), previously equilibrated with buffer containing 50 mM Tris pH 7.5, 400 mM NaCl, 10% glycerol, and 5 mM β-mercaptoethanol or 1 mM DTT. Fractions containing AsCas12a were identified by SDS-PAGE using 10% acrylamide gels and pooled. The purified protein was sterile-filtered through a 0.2 μm filter, flash-frozen in liquid nitrogen, and stored in aliquots at −80 °C. Protein concentration was determined by measuring absorbance at 280 nm using a NanoDrop One spectrophotometer (Thermo Fisher Scientific) and applying a molar extinction coefficient of 148,410 M−1 cm−1. All purification steps were performed at 4 °C.

To generate an MBP-tagged SpCas9 protein optimized for expression in *E. coli*, the NsiI restriction site was first introduced at the C-terminus of the MBP sequence in the MBP-SpCas9 plasmid (Addgene plasmid #88915) by site-directed mutagenesis. The resulting plasmid was digested with NsiI and BstEII to enable the subcloning of the *E. coli* codon optimized Cas9 sequence, which was amplified from the SpCas9-Ec plasmid (Addgene plasmid #62731). For SpCas9 purification, the newly generated Cas9-2NLS bacterial expression plasmid was transformed into competent *E. coli* BL21 cells. SpCas9 expression and purification were subsequently performed using the same procedures described above for AsCas12a.

### 4.2. MNP-RNP nanocomplexes characterization

#### 4.1.1. MNP synthesis and functionalization

The γ-Fe2O3 (maghemite) cores were prepared using a co-precipitation procedure combined with an acid treatment. Briefly, 24.3 g of FeCl3·6H2O (Sigma-Aldrich) and 24.3 g of FeCl2·4H2O (Sigma-Aldrich) were fully dissolved in 0.425 L of ultrapure water at room temperature under continuous stirring.

Next, 25% NH4OH (sigma-Aldrich) was added at a rate of 0.1 mL/s and stirred for 5 more minutes. The temperature was then raised to 90°C for 3 hours. Finally, the mix was incubated at room temperature overnight. After overnight incubation, magnetic decantation was used to isolate the resulting precipitate. To eliminate the excess of iron from the NP surface, the precipitate was treated with 300 mL of 2M HNO3 (Sigma-Aldrich) for 15 minutes. Again, magnetic separation was used to remove HNO3. Next, 130 mL of water and 75 mL of 1M Fe (NO3)3·9H2O 1M (sigma-Aldrich) were added to the precipitate. The mixture was boiled for 30 minutes under continuous shaking and subsequently cooled down. Finally, the mixture was treated with 300 mL of 2M HNO3 for 15 minutes. The resulting γ-Fe2O3 cores were wahed with water sveral times. A rotary evaporator was used to concentrate the cores.

The maghemite cores were next coated with carboxymethyldextran (CMD) by treating them with an 80 mg/mL solution of CMD (Sigma-Aldrich) in water. Next, 65% HNO3 was added to lower the pH to 3.0 and the mixture was sonicated for 10 hours. After sonication, the pH was raised to 7.0.

Particle morphology and size distribution were characterized using a JEOL JEM 1010 transmission electron microscope (TEM). A single drop of the diluted sample suspension was deposited onto a carbon-coated copper grid and allowed to dry overnight under ambient conditions. Particle shape and size distribution were determined by manual analysis of over 1400 individual particles using ImageJ Software (Fiji).

#### 4.1.2. MNP-RNP Cas nucleases interaction

MNPs were pelleted by centrifugation (13200 rpm, 45 minutes, 4°C), resuspended to a final concentration of 6 mg/mL and dispersed by sonication for 5 minutes in an ultrasonic bath. Cas9 and Cas12a RNPs were assembled by mixing the Cas nuclease at a final concentration of 9.6 µM with the corresponding crRNA (Table S2) in a 1:1 molar ratio, followed by incubation at room temperature for 15 minutes. Next, RNPs (2.4 µM final concentration) and MNPs (3 mg/mL final concentration) were mixed in interaction buffer containing 50 mM Tris pH 7.5, 100 mM NaCl and 10% glycerol. This mixture was incubated for 1 h at room temperature, followed by 4 hours at or overnight at 4°C, in a rotating well to ensure sample homogeneity.

#### 4.1.3. Evaluation of MNP-RNP interaction

To pellet MNP-RNP complexes and remove unbound protein, samples were centrifuged at 8000 rpm for 15 minutes at 4°C. The supernatant (unbound protein fraction) was collected and stored, while the pellet was resuspended in interaction buffer (50 mM Tris pH 7.5, 100 mM NaCl and 10% glycerol). Both fractions were analysed by SDS-PAGE using Mini-PROTEAN TGX Stain-Free Gels (Bio-Rad). ImageJ Software (Fiji) was used to quantify binding efficiency by gel densitometry according to predefined criteria. In general, binding was calculated based on the pellet fraction (Equation 1). However, in some cases, the protein bound to the MNPs remained associated to the nanoparticles during the SDS-PAGE, despite the presence of SDS. This resulted in a lower band intensity in the pellet fraction, and consequently the sum of the supernatant and pellet band intensities in the MNP-RNP samples was lower than that of the RNP alone control. In these cases, binding was instead calculated from the supernatant fraction, corresponding to the protein remaining unbound, and was expressed relative to the total protein in the RNP-alone control (Equation 2).

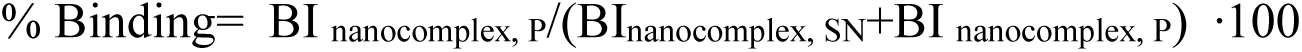

Equation 1: Estimation of the percentage of protein bound to MNPs based on the pellet fraction. BI: band intensity, SN: supernatant, P: pellet.

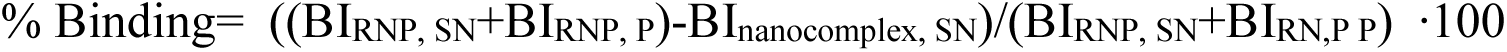

Equation 2: Estimation of the percentage of protein bound to MNPs based on the supernatant fraction. BI: band intensity, SN: supernatant, P: pellet.

#### 4.1.4. MNP-RNP characterization using light scattering

Nanoconjugates’ hydrodynamic size and zeta-potential were measured using Zetasizer Nano ZS (Malvern Panalytical). Samples were diluted to 0.1 mg Fe (mL in interaction buffer (50 mM Tris pH 7.5, 100 mM NaCl and 10% glycerol). Size measurements were performed in 70 µL disposable microcuvettes (Brand). Z-potential was assessed using folded capillary Zeta cells (Malvern Panalytical). All measurements were carried out at 25°C.

#### 4.1.5. MNP-RNP nanocomplex stability in biological media

To study the colloidal stability and the stability of the MNP-RNP interaction, MNP-RNPs were incubated in Dulbecco’s Modified Eagle Medium (DMEM, Thermofisher Scientific) supplemented with 10% fetal bovine serum (FBS, Thermofisher Scientific) and 1% streptomycin-penicillin (Thermofisher Scientific) at room temperature for 4 or 24 hours. MNP-RNPs incubated in interaction buffer (50 mM Tris pH 7.5, 100 mM NaCl and 10% glycerol) were used as a control. The hydrodynamic size and the percentage of bound protein were measured according to the protocols described above.

### 4.2. Cell lines

For in vitro studies, HEK-293-uGFP were purchased from Amsbio (SC058). HEK-293T cell line was gifted by Dr. Sandra Rodríguez (CNIO), MEF-tdTomato cells were obtained from Dr. Mirentxu Santos (CIEMAT) and HDFn cells were kindly provided by Dr. Marta García (CIEMAT). All cell lines and primary cells were cultured in Dulbecco’s Modified Eagle Medium (DMEM, Thermofisher Scientific) supplemented with 10% fetal bovine serum (FBS, Thermofisher Scientific) and 1% streptomycin-penicillin (Thermofisher Scientific). Cells were cultured at 37°C and 5% CO2.

### 4.3. Treatment of cell cultures with nanocomplexes

Cells were seeded 24h prior to in vitro experiments in 96-well plates and treated at 70-80% confluency. For cell treatments, the culture medium was replaced with fresh medium containing MNP-RNP nanocomplexes at a final concentration of 1 mg/mL MNPs and 800 nM RNPs, unless otherwise indicated. Incubation times varied by cell line: HEK-293-uGFP and MEF-tdTomato cells were incubated for 4 hours at 37°C; whereas HeLa, HEK-293T and HDFn cells were incubated overnight. Following incubation, the nanocomplex-containing medium was removed and replaced with fresh growth medium. The cells were then maintained at 37 °C and 5% CO2, and subsequent analyses were performed at the specified time points.

### 4.4. Cellular uptake

#### 4.4.1. Prussian Blue staining

After O/N incubation with MNP-RNPs, cells were first washed 3 times with PBS and next fixed for 5 minutes with cold methanol. Coverslips were allowed to air-dry before proceeding with the staining. Iron was stained with a 1:1 mix of 4% potassium hexacyanoferrate trihydrate and 4% HCl for 15 minutes at room temperature. After washing 3 times with PBS, cells were counterstained with 0.02% of neutral red. The excess of neutral red was eliminated by immersing the coverslips in distilled water. Samples were imaged with Leica DM2000 LED (Leica Microsystems) optical microscope.

#### 4.4.2. MNP-RNP cellular internalization

ATTO-labelled Cas RNPs were used to form MNP-RNP nanocomplexes following the same procedure as for unlabeled proteins, but at 2X concentration. 12,000 HeLa cells were seeded per well in 96-well plates with optical bottom (Ibidi) and treated with MNP-RNP complexes. Final RNP concentration in the well was 50 nM. After the 3-hour incubation, media was aspirated and replaced by new media containing Hoechst (1:10000, Thermofisher Scientific) and SPY555-actin (1:1000, Spyrochrome). Cells were imaged 6-8 hours after treatment using live-cell confocal laser scanning microscopy (SP-5, Leica).

### 4.5. Cellular effects of MNP-RNPs

#### 4.5.1. Cell death

HEK-293T cells were treated with MNP-RNP nanocomplexes for 24 hours. 5% DMSO was used as a positive biological control. Apoptotic and necroptotic cells were evaluated at 24, 48 and 72 hours. At each time point, supernatants were collected and cells were trypsinized. The obtained cells were washed with PBA, centrifuged at 1400 rpm for 5 minutes and labeled with FITC Annexin V (BD Pharmingen) and Propidium Iodide (SIGMA) 1 µg/UI diluted in Annexin Binding Buffer 1X (BD Pharmingen). Cells were incubated for 15 minutes in darkness at room temperature. After that, 300 µL Annexin Binding Buffer 1X were added and data was acquired using CytoFlex Flow Cytometer (Beckman Coulter).

#### 4.5.2. Cell cycle

HEK-293T cells were treated with MNP-RNP nanocomplexes for 24 hours and Colchicine at 1 µg/mL was used as biological control. Cell cycle distribution was evaluated at 24 and 48 hours. At each time point, supernatants were collected and cells were trypsinized. The obtained cells were washed with PBS 1X and fixed by adding 200 µL of cold 70% ethanol. Samples were incubated at 4°C for fixation for at least 2 hours. Following fixation, cells were centrifuged at 1400 rpm for 7 minutes, resuspended in 200 µL of PBS 1X supplemented with 50 µg/mL RNase A and incubated for 20 minutes at 37°C. Propidium Iodide (SIGMA) was subsequently added to a final concentration of 50 µg/mL and cells were incubated for 15 minutes in darkness at room temperature. Data acquisition was performed using CytoFlex Flow Cytometer (Beckman Coulter).

### 4.6. Gene editing evaluation

#### 4.6.1 Reporter cell lines flow cytometry

5 days after transduction, cells’ fluorescence was directly analyzed by flow cytometry using CytoFlex Flow Cytometer (Beckman Coulter). The percentage of GFP-negative cells and tdTomato-positive cells were then quantified. The efficiency of gene editing by MNP-RNPs was compared to the commercial Lipofectamine-2000 reagent (ThermoFisher).

#### 4.6.2. B2M flow cytometry

B2M knock out was evaluated 5 days after transduction. 2·105 transduced cells were washed with PBA, centrifuged at 1,400 rpm for 5 minutes, and stained with PE HLA-ABC antibody (BioLegend) for 20 minutes at 4°C in the dark. Following an additional wash with PBA, the cells were counterstained with DAPI and analyzed by flow cytometry using CytoFlex Flow Cytometer (Beckman Coulter). Cell population gates were manually defined using FlowJo Software (BD) and the percentage of B2M-negative cells were then quantified.

#### 4.6.3. Digital droplet PCR (ddPCR)

Genomic DNA was extracted using DNeasy Blood & Tissue Kit (Qiagen). A 1X digital PCR supermix for probes (No dUTP) (Bio-Rad) was mixed with an assay mixture containing primers (0.9 µM, Table S3) and probes (0.25 µM, Table S3). The reaction mixture was distributed at equal volumes into a ddPCR 96-well plate (Bio-Rad). Hind-III (NEB) digested DNA (100 ng per assay) was then added, mixed by vortexing, and the plate was sealed with a PCR Plate Heat Seal (Bio-Rad).

Droplets were generated in a new 96-well plate using Automated Droplet Generation Oil for Probes (Bio-Rad) and DG32 Automated Droplet Generator Cetridges (Bio-Rad) in an automated droplet generator (Bio-Rad), after which the plate was sealed. PCR amplification was performed using C1000 Touch Thermal Cycler (Bio-Rad), and the droplets were analyzed with a QX200 Droplet Reader (Bio-Rad). Droplet thresholds were manually stablished using CX Manager Software (Bio-Rad), and the percentage of NHEJ events was the quantified.

#### 4.6.4. Polymerase Chain Reaction (PCR) to assess Col7A1 exon 80 deletion

To assess Col7A1 exon 80 deletion in HDFn cells, genomic DNA was extracted using DNeasy Blood & Tissue Kit (Qiagen). PCR amplification of the targeted Col7A1 gene region was performed using FastStart Taq DNA Polymerase (Roche). Reactions were prepared in a total volume of 15 µL containing FastStart PCR Buffer 1X (with 2 mM MgCl2), dNTP mix (0.17 µM), specific primers (1.3 µM, Table S3), 1.25 U FastStart Taq DNA Polymerase, and 3 µL of DNA template. PCR program touchdown 65-60°C was as follows: 94 °C for 4 min; 5 cycles of 94°C for 30 s, 65°C for 30 s and 72°C por 30 s, decreasing annealing temperature 1°C every cycle; followed by 30 cycles of 94 °C for 30 s, 60°C for 30 s, and 72 °C for 30 s; then 72 °C for 7 min. PCR products were resolved on a 2% agarose gel in 1X TAE buffer containing GeenSafe Premium (NZYtech). Molecular wight marker was 100 bp DNA Ladder RTU (Bio-Helix).

## 6. Author Contributions

JPM: formal analysis, investigation, methodology, visualization, writing-original draft. CEM: conceptualization, investigation, methodology, visualization. MD: investigation, resources. NLG: investigation, resources, methodology. MC: Resources. CMC: resources. Yurena Luengo: resources, methodology. GS: resources, supervision. MG: resources, methodology. MS: resources, methodology. AM: resources, methodology. RM: resources, methodology. AS: resources, supervision, funding acquisition. BS: supervision, funding acquisition, conceptualization, project administration, methodology, writing-original draft, writing-review and editing.

