## Supplementary material for "Carboxymethyl Dextran Coated Iron Oxide Magnetic Nanoparticles as Vehicles for CRISPR Ribonucleoproteins": Suplemental figures

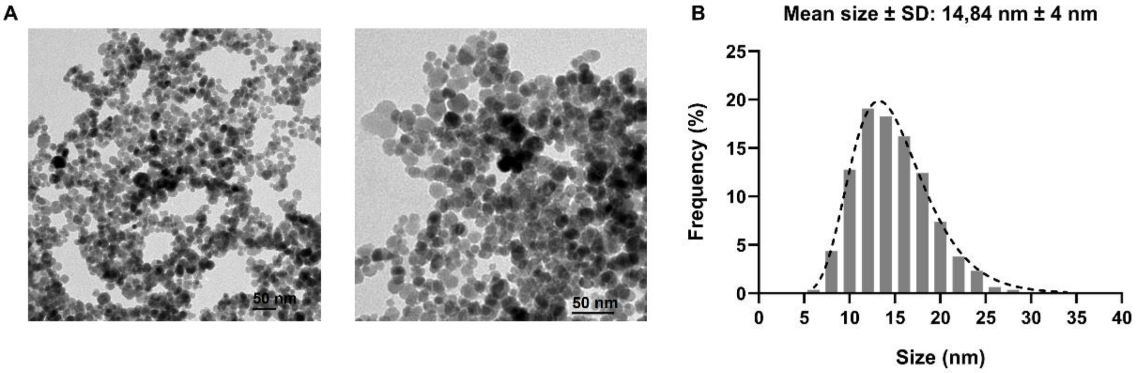


**Figure S1**. A) TEM micrographs. Scale bar: 50 nm. B) Size distribution of 14 nm $Fe_2O_3$ cores of uncoated MNPs.

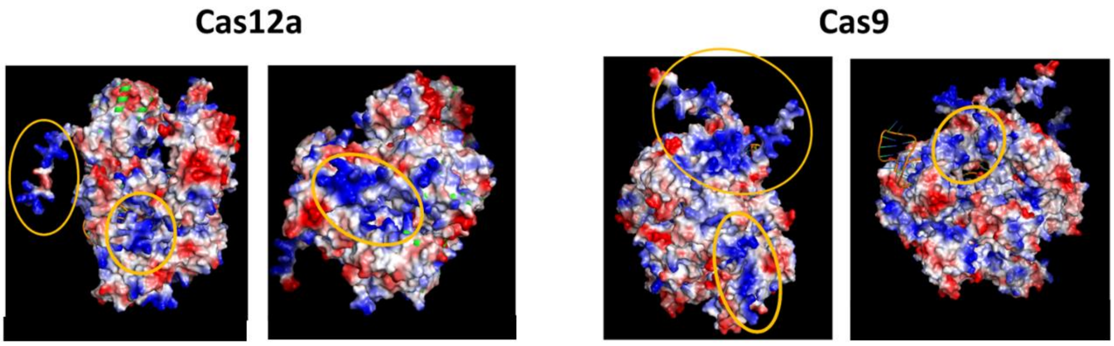


**Figure S2**. Electrostatic charge of Cas9 and Cas12a RNPs surface visualized by pyMOL. The structure of AsCas12a-HAepitope-NLS-NLS and NLS-Cas9-NLS-NLS was determined by AlphaFold. Positive surfaces were labeled by orange circles.

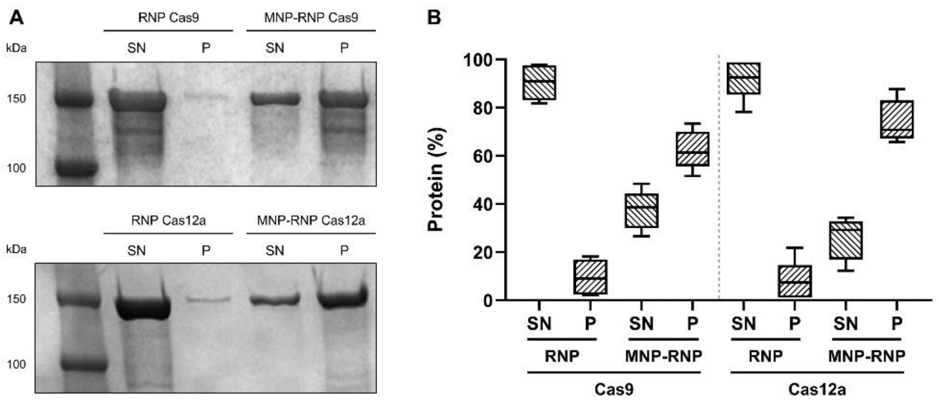


**Figure S3**. Interaction between MNPs and Cas RNPS. A) MNP-RNPs interaction was analyzed by PAGE-SDS and subsequent densitometry. B) Percentage of protein detected in RNPs alone or complexed with MNPs. SN: supernatant, P: pellet. (n=5).

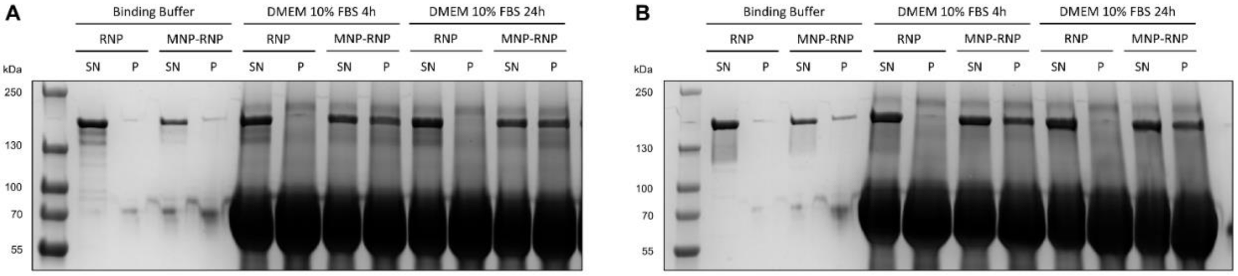


**Figure S4**. SDS-PAGE gel showing interaction stability of A) Cas9 RNPs and B) Cas12a RNPs complexed with MNPs after 4 or 24h of incubation in DMEM supplemented with 10% FBS. Interaction was quantified by densitometry. SN: supernatant, P: pellet.

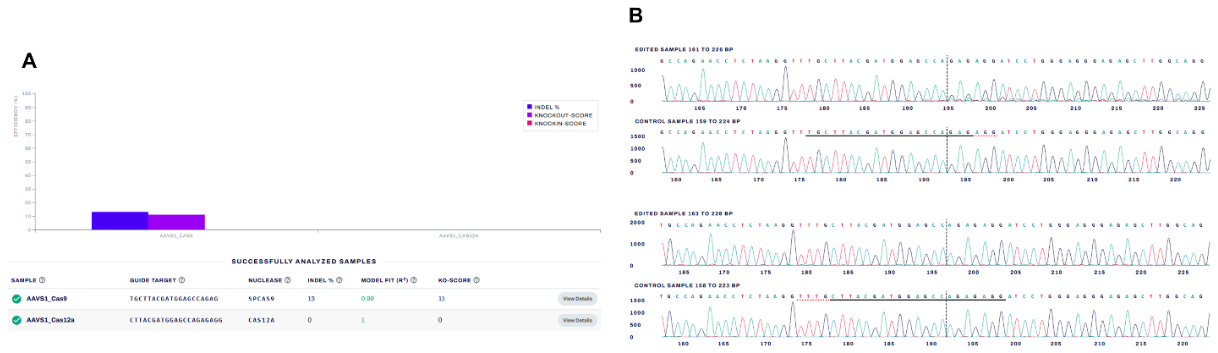


**Figure S5.** Sanger sequencing analysis using EditCo's Interference of CRISPR Edits (ICE) Tool. A) Percentage of indels and knockout score for Cas9 and Cas12a target sites in HEK-293T cells. B) Chromatogram trace files comparing edited sample against the untreated control sample.

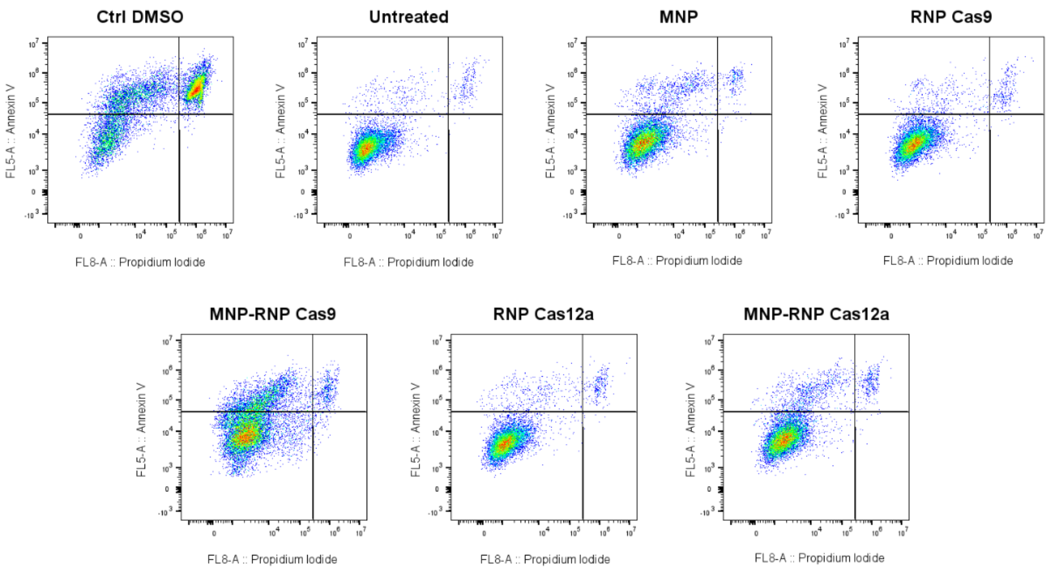


**Figure S6**. Representative flow cytometry plots showing Annexin V fluorescence intensity versus Propidium Iodide fluorescence intensity staining in HEK-293T cells 24 hours after treatment. Conditions tested include biological DMSO control (positive control for apoptosis/necrosis), untreated cells, naked MNPs, uncomplexed RNPs and MNP-RNP nanocomplexes. Quadrant delineate the different cell populations: viable cells (lower-left), necrotic cells (lower-right), early-apoptotic cells (upper-left) and late apoptotic cells (upper-right).

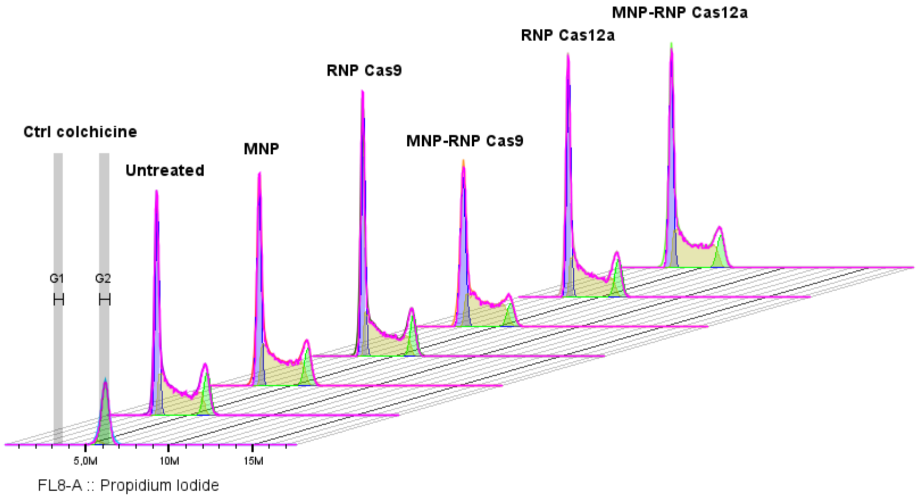


**Figure S7**. Representative flow cytometry offset histogram profiles showing cell cycle distribution based on Propidium Iodide DNA staining in HEK-293T cells 24 hours after treatment. Conditions tested include biological Colchicine control (positive control for G2/M phase arrest), untreated cells, naked MNPs, uncomplexed RNPs and MNP-RNP nanocomplexes. Histogram peaks and fitted curves delineate the specific cell cycle phases: G0/G1 phase (first major peak), S phase (intermediate region), and G2/M phase (second peak).

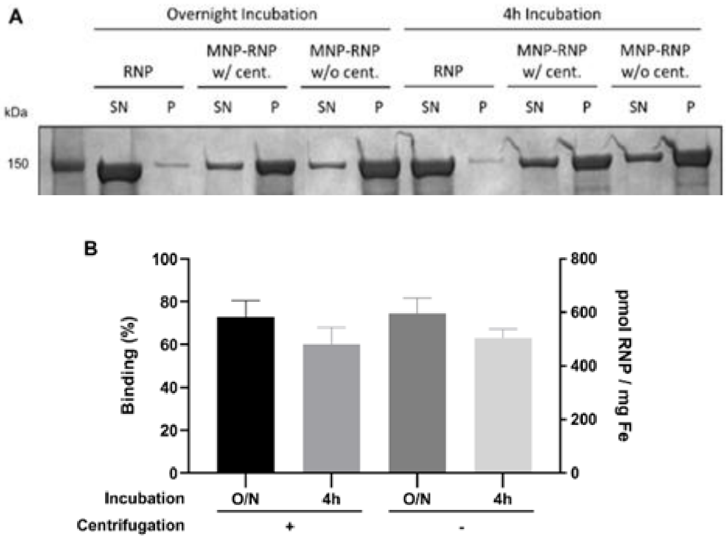


**Figure S8**. A) SDS-PAGE gel analysis evaluating the binding efficiency of MNP-RNP under different assembly conditions: incubation time (Overnight versus 4h) and centrifugal processing (with centrifugation versus without centrifugation). SN: supernatant, P: pellet. B) Quantification of binding efficiency by densitometry (n=3).

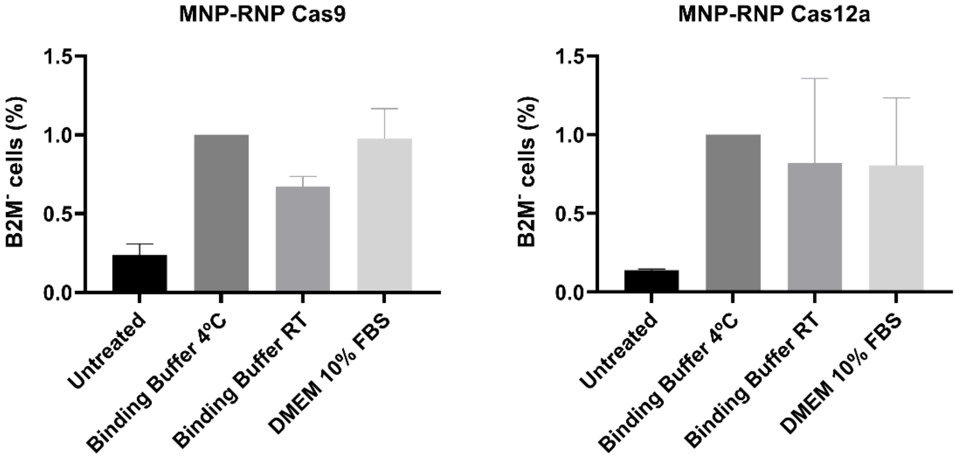


**Figure S9.** Gene editing efficiency of nanocomplexes in HEK-293T cells following incubation in biological media. Nanocomplexes were incubated for 4 hours in DMEM supplemented with 10% FBS at room temperature (RT), whereas control samples in Binding Buffer were maintained at RT or 4ºC. Data were normalized to the MNP-RNP control in Binding Buffer at 4ºC. Percentage of B2M knockout cells evaluated by flow cytometry.

**Table S1**. Hydrodynamic size (diameter size, PDI: polydispersity index) of uncoated MNPs, MNP-RNP Cas9 and MNP-RNP Cas12a after 0, 4 and 24 hours of incubation in DMEM supplemented with 10% FBS. Mean ± SD, n=3.

| *Sample* | *Time point* | *Size (nm) ± SD* | *PDI ± SD* |
|---|---|---|---|
| **MNP** | 0h | 250.8 ± 8.30 | 0.76 ± 0.11 |
| | 4h | 364.6 ± 15.17 | 0.62 ± 0.12 |
| | 24h | 314.5 ± 12.75 | 0.47 ± 0.00 |
| **MNP-RNP Cas9** | 0h | 117.3 ± 0.24 | 0.21 ± 0.02 |
| | 4h | 142.6 ± 1.40 | 0.23 ± 0.02 |
| | 24h | 158.2 ± 0.96 | 0.19 ± 0.00 |
| **MNP-RNP Cas12a** | 0h | 118.0 ± 0.91 | 0.21 ± 0.01 |
| | 4h | 146.3 ± 0.35 | 0.22 ± 0.01 |
| | 24h | 164.2 ± 1.08 | 0.20 ± 0.01 |

**Table S2**. sgRNAs for Cas9 and Cas12a-based KO.

| *sgRNA Name* | *Target Sequence (5'-3')* |
|---|---|
| **Cas12a HEK-GFP** | CGTCGCCGTCCAGCTCGACCA |
| **Cas9 MEF-tdTomato** | TCCTATTCTCTAGAAAGTAT |
| **AAVS1_Cas9** | TGCTTACGATGGAGCCAGAG |
| **AAVS1_Cas12a** | CTTACGATGGAGCCAGAGAGG |
| **B2M_Cas9** | GAGTAGCGCGAGCACAGCTA |
| **B2M_Cas12a** | AGTGGGGGTGAATTCAGTGT |
| **COL7A1 Exon80 sg2** | CTCTATGTAGGGTCTGCAGG |
| **COL7A1 Exon80 sg3** | AAGAACCTTCACCTGTC |

**Table S3**. Amplification primers and probes for gene editing detection.

| *Use* | *Target* | *Sequence (5'-3')* |
|---|---|---|
| **Primers for NHEJ analysis by ddPCR** | B2M_Cas12a Forward | CCATCCGACATTGAAGTTGAC |
| | B2M_Cas12a Reverse | CACTTAACTATCTTGGGCTGTG |
| | B2M_Cas9 Forward | TTGTCCTGATTGGCTGG |
| | B2M_Cas9 Reverse | GCGGGAGAGGAAGGA |
| **Probes for NHEJ analysis by ddPCR** | B2M_Cas12a WT | 56-FAM/AGTGG+AGCATTCA+GA+CTTG/3IABkFQ/ |
| | B2M_Cas12a NHEJ | 5HEX/TAT+CTCTTGTA+CTA+CA+CTGA/3IABkFQ/ |
| | B2M_Cas9 WT | /56-FAM/TATAAGTG+GAGGC+GT+CGTC/3IABkFQ/ |
| | B2M_Cas9 NHEJ | 5HEX/C+GAG+CACA+G+CTAAGG/3IABkFQ/ |
| **COL7A1 Exon80 PCR** | COL7A1 Exon80 Forward | AGGGGTCAGGACGGCAAC |
| | COL7A1 Exon80 Reverse | CAGCTCCAGTAGGTCCAGTCAG |
